# Draft genome assembly and population genetics of an agricultural pollinator, the solitary alkali bee (Halictidae: *Nomia melanderi*)

**DOI:** 10.1101/465351

**Authors:** Karen M. Kapheim, Hailin Pan, Cai Li, Charles Blatti, Brock A. Harpur, Panagiotis Ioannidis, Beryl M. Jones, Clement F. Kent, Livio Ruzzante, Laura Sloofman, Eckart Stolle, Robert M. Waterhouse, Amro Zayed, Guojie Zhang, William T. Wcislo

## Abstract

Alkali bees (*Nomia melanderi*) are solitary relatives of the halictine bees, which have become an important model for the evolution of social behavior, but for which few solitary comparisons exist. These ground-nesting bees defend their developing offspring against pathogens and predators, and thus exhibit some of the key traits that preceded insect sociality. Alkali bees are also efficient native pollinators of alfalfa seed, which is a crop of major economic value in the United States. We sequenced, assembled, and annotated a high-quality draft genome of 299.6 Mbp for this species. Repetitive content makes up more than one-third of this genome, and previously uncharacterized transposable elements are the most abundant type of repetitive DNA. We predicted 10,847 protein coding genes, and identify 479 of these undergoing positive directional selection with the use of population genetic analysis based on low-coverage whole genome sequencing of 19 individuals. We found evidence of recent population bottlenecks, but no significant evidence of population structure. We also identify 45 genes enriched for protein translation and folding, transcriptional regulation, and triglyceride metabolism evolving slower in alkali bees compared to other halictid bees. These resources will be useful for future studies of bee comparative genomics and pollinator health research.

## Introduction

The comparative method is required for sociogenomics research, which aims to explain how social behavior evolves from a molecular perspective within the context of Darwinian evolution (Robinson *et al*. 2005). Eusociality is a special form of social behavior in animals that involves extreme levels of cooperation at the level of the group, manifest as queens and workers who distribute tasks related to reproduction, brood care, nest maintenance, and defense within a colony (Wilson 1971). A large amount of comparative genomics research has focused on the insect order Hymenoptera, because ants, bees, and wasps display remarkable variation in social organization, and they represent at least five independent origins of eusociality in the past 200 million years (Danforth *et al*. 2013; Branstetter *et al*. 2017). The comparative method is most powerful for understanding social evolution when it includes closely related species that are representative of the solitary ancestor from which eusociality arose (Rehan and Toth 2015). However, the rate at which genomic resources have become available for social Hymenoptera has far out-paced that for solitary species. Genome assemblies are publicly available for just three solitary bees and no solitary vespid wasps, compared to over 30 reference genomes currently available for social bees, wasps, and ants (Branstetter *et al*. 2018). This is in stark disproportion to the species that express solitary behavior among bees and wasps, most of which lead solitary lifestyles (Wcislo and Fewell 2017).

Alkali bees (*Nomia melanderi*) belong to the subfamily Nomiinae (Halictidae), a taxon composed of species that are solitary, though some express communal behavior and other forms of social tolerance (Wcislo and Engel 1996). The subfamily is the sister clade to the Halictinae, which includes both solitary and social lineages (Danforth *et al*. 2008). The alkali bees may be representative of the solitary ancestor from which eusociality likely evolved within the bee family Halictidae, and provide important phylogenetic context to comparative genomics (Brady *et al*. 2006; Gibbs *et al*. 2012). Alkali bees also possess several of the characteristic traits thought to be important in the ancestor of social halictids, including nest defense and other forms of maternal care (Batra and Bohart 1969; Batra 1970, 1972) (Fig. 1A). As such, this species has become an important model for testing hypotheses for the origins of eusociality, and has provided meaningful insight into the reproductive physiology of solitary bees (Kapheim 2017; Kapheim and Johnson 2017a, 2017b). Development of genomic resources for this species will enable additional hypothesis testing regarding the solitary antecedents of eusociality in this family, and insects in general.

**Figure 1.**
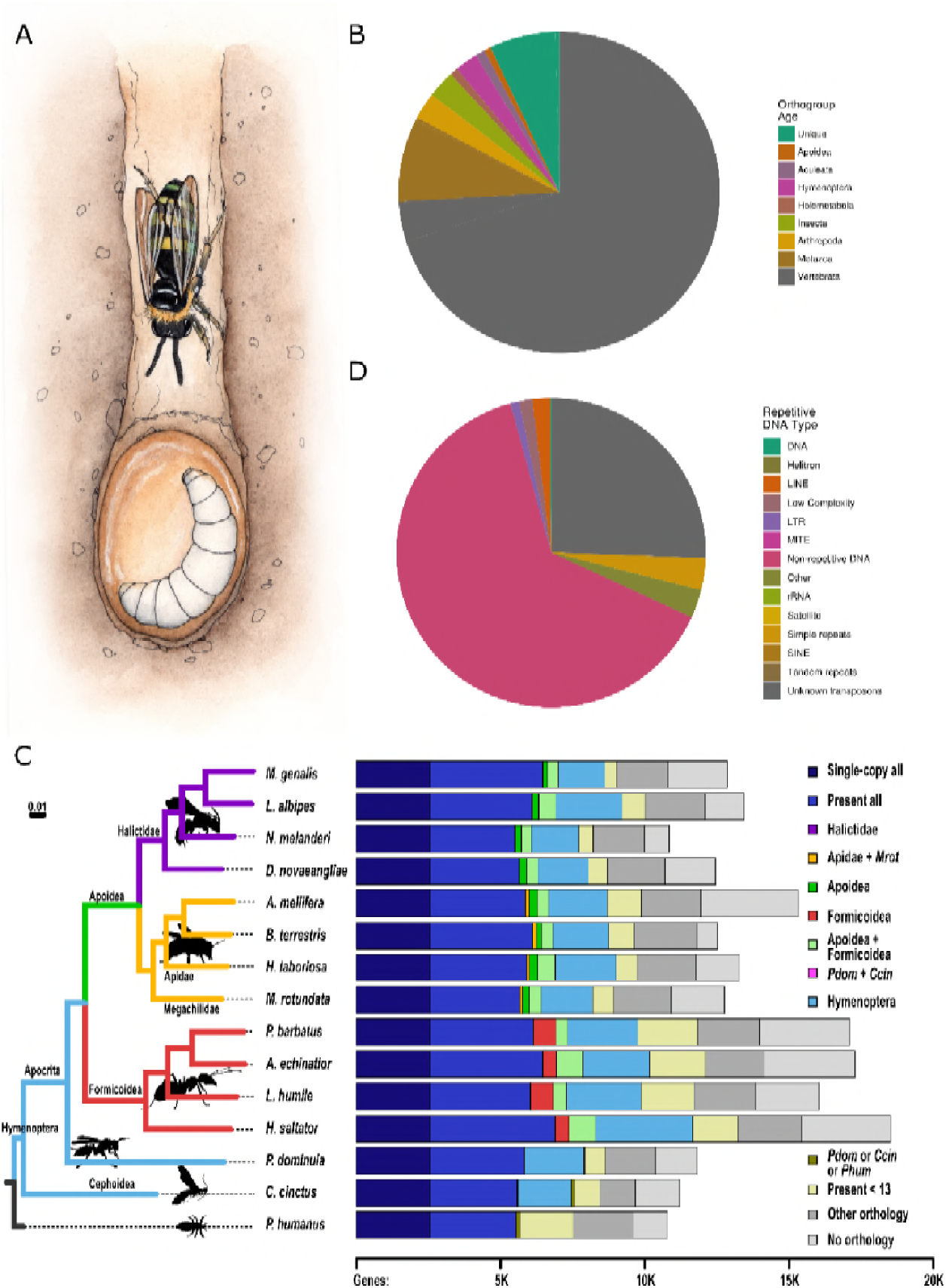
*Nomia melanderi* genome characteristics and comparative context. (A) *N. melanderi* are ground-nesting bees with maternal care. (B) Most of the protein-coding genes belong to OGs that include vertebrates or other metazoans, and are thus widely conserved. (C) *N. melanderi* species phylogeny (left) and gene orthology (right). The maximum likelihood 15-species molecular phylogeny estimated from the superalignment of 2,025 single-copy orthologs recovers supported families. Branch lengths represent substitutions per site, all nodes achieved 100% bootstrap support. Right: Total gene counts per species partitioned into categories from single-copy orthologs in all 15 species, or present but not necessarily single-copy in all (i.e., including gene duplications), to lineage-restricted orthologs (Halictidae, Apidae and *M. rotundata*, Apoidea, Formicoidea, Apoidea and Formicoidea, Hymenoptera, specific outgroups), genes showing orthology in less than 13 species (i.e., patchy distributions), genes present in the outgroups (present in *P. domunila* or *C. cinctus*, present in *P. dominula* or *C. Cinctus* or *P. humanus*), and genes with orthologs from other sequenced insect genomes or with no identifiable orthology. The purple Halictidae bar is present but barely visible as only 16 to 32 orthologous genes were assigned to the Halictidae-restricted category. (D) A large proportion of repetitive DNA consists of uncharacterized transposable elements, but all major transposon groups were detected.

The development of genomic resources for alkali bees will also have practical and applied benefits. Alkali bees are native pollinators of alfalfa seed, which is a multi-billion dollar industry in the United States, accounting for one-third of the $14 billion value attributed to U.S. bee-pollinated crops (Van Deynze *et al*. 2008; U.S. Department of Agriculture 2014). With issues of honey bee health and colony loss over the last decade, increased attention has been placed on the need to find alternative pollinators for many of our most important crops. Aggregations of alkali bees have been sustainably managed alongside alfalfa fields in southeastern Washington state for several decades (Cane 2008), and they are more effective pollinators of this crop than honey bees (Batra 1976; Cane 2002). Moreover, as a naturally aggregating native species, they are less costly pollinators than alfalfa leafcutter bees (*Megachile rotundata*), which must be purchased commercially (James 2011). Genomic resources have been an invaluable resource for the study of honey bee health and management, and are thus likely to benefit this important pollinator as well.

Here we present a draft genome assembly and annotation for *N. melanderi*, along with intial genomic comparisons with other Hymenoptera, a description of transcription factor binding sites, and population genetic analyses based on resequencing of individuals from throughout the southeastern Washington population. These resources will provide an important foundation for future research in sociogenomics and pollinator health.

## Materials and Methods

### Genome sequencing and assembly

#### Sample collections

All of the bees used for sequencing were collected from nesting aggregations in and around Touchet, Washington (USA) with permission from private land owners in June 2014 or June 2015. Adult males and females were captured live, and flash frozen in liquid nitrogen. They were transported in a dry nitrogen shipper, and then stored at -80 ^°^C until nucleic acid extraction.

#### DNA and RNA isolation

For genome sequencing, we isolated genomic DNA from individual males in three separate reactions targeting either the head or one half of a thorax. We used a Qiagen MagAttract kit, following the manufacturer’s protocol, with two 200 μl elutions in AE buffer. We isolated RNA from three adult females using a Qiagen RNeasy kit, following the manufacturer’s protocol, eluting once in 50 μl of water. We extracted RNA from the head and rest of the body separately for each female. For whole genome resequencing, we isolated genomic DNA of 18 adult females and one male from half of a thorax with a Qiagen MagAttract kit, as above. DNA was quantified with a dsDNA high sensitivity Qubit reaction, and quality was assessed on an agarose gel. RNA was quantified on a Nanodrop spectrophotometer, and quality was assessed with a Bioanalyzer.

#### Sequencing

All library preparation and sequencing was performed at the Roy J. Carver Biotechnology Center at University of Illinois at Urbana-Champaign. Two shotgun libraries (350-450 bp, 500-700 bp) were prepared from the DNA of a single haploid male with the Hyper Kapa Library Preparation kit (Kapa Biosystems). Three mate-pair libraries (3-5 kb, 8-10 kb, 15-20 kb) were constructed from DNA pooled from five individual males using the Nextera Mate Pair Library Sample Prep kit (Illumina, CA), followed by the TruSeq DNA Sample Prep kit. A single RNA library was constructed from pooled RNA from the six female tissue samples with the TruSeq Stranded mRNA Library Construction kit (Illumina, CA).

DNA libraries were quantitated by qPCR and sequenced on a HiSeq2500 for 251 cycles from each end of the fragments using a TruSeq Rapid SBS kit version 2. Shotgun libraries were sequenced on a single lane, and mate-pair libraries were pooled and sequenced on a single lane. RNA libraries were sequenced on a single lane for 161 cycles from each end of the fragments. Fastq files were generated and demultiplexed with the bcl2fastq v1.8.4 Conversion Software (Illumina).

#### Genome assembly

The DNA shotgun and mate-pair library sequencing generated a total of 593,526,700 reads. After adapter trimming, these reads were filtered for quality (Phred 64 < 7) and excessive (≥10) Ns. We removed PCR duplicates from read pairs.

We used SOAPdenovo 2 with default parameters for genome assembly. We began by constructing contigs from the shotgun library reads split into kmers, which were used to construct a de Bruijn graph. Filtered reads were then realigned onto the contigs, and used to construct scaffolds based on shared paired-end relationships between contigs. We then closed gaps in the assembly using information from paired-end reads that mapped to a unique contig and a gap region.

#### BUSCO assessment of assembly completeness

The genome assembly completeness in terms of expected gene content was quantified using the Benchmarking Universal Single-Copy Ortholog (BUSCO) assessment tool (Waterhouse *et al*. 2018) for *N. melanderi* and seven other Apoidea species. Assembly completeness assessments employed BUSCOv3.0.3 with Augustus 3.3 (Stanke *et al*. 2006), HMMER 3.1b2 (Finn *et al*. 2011), and BLAST+ 2.7.1 (Camacho *et al*. 2009) (Camacho et al. 2009), using both the hymenoptera_odb9 and the insecta_odb9 BUSCO lineage datasets and the Augustus species parameter ‘honeybee1’.

### Genome annotation

#### Gene annotation

We predicted gene models based on homology and *de novo* methods. Results were integrated with GLEAN (Elsik *et al*. 2014). Homology based gene prediction used the gene models of four species (*Apis mellifera*, *Acromyrmex echinator*, *Drosophila melanogaster*, and *Homo sapiens*). We used TBLASTN to gather a non-redundant set of protein sequences, and then selected the most similar proteins for each candidate protein coding region based on sequence similarity. Short fragments were connected with a custom script (SOLAR), and Genewise (v2.0) (Birney *et al*. 2004) was used to generate the gene structures based on the homology alignments. This generated four gene sets, based on homology with four different species.

We used Augustus (Stanke *et al*. 2006) and SNAP (Johnson *et al*. 2008) for *de novo* gene prediction, with parameters trained on 500-1,000 intact genes from the homology-based predictions. We chose genes that were predicted by both programs for the final *de novo* gene set.

The four homology-based gene sets and one *de novo* gene set were integrated to generate a consensus gene set with GLEAN. We then filtered genes affiliated with repetitive DNA and genes whose CDS regions contained more than 30% Ns. Repetitive DNA was identified through annotation of tandem repeats (Tandem Repeats Finder v4.04) (Benson 1999) and transposable elements (TEs). This initial identification of TEs was performed based on homology-based and *de novo* predictions. For the homology-based approach, we used RepeatMasker (v3.2.9) and RepeatProteinMask (v3.2.9) (“Smit AFA, Hubley R, Green P: RepeatMasker. Available at: http://www.repeatmasker.org. [Accessed 9 April 2013]”) against a custom build of the Repbase library. *De novo* predictions were performed with LTR_FINDER (v1.0.5) (Xu and Wang 2007), PILER (v1.0) (Edgar and Myers 2005), and RepeatScout (v1.0.5) (Price *et al*. 2005). Results were used as an input library for a second run of RepeatMasker.

We used the 571,457,212 reads generated from RNA sequencing to polish the gene set. After filtering, we mapped reads to the genome with TopHat (Trapnell *et al*. 2009), and used Cufflinks (Trapnell *et al*. 2012) to assemble transcripts. Assembled transcripts were then used to predict ORFs. Transcript-based gene models with intact ORFs that had no overlap with the GLEAN gene set were added. GLEAN gene models were replaced by transcript-based gene models with intact ORFs when there was a discrepancy in length or merging of gene models. Transcripts without intact ORFs were used to extend the incomplete GLEAN gene models to find start and stop codons.

Putative gene functions were assigned to genes based on best alignments to the Swiss-Prot database (Release 2013_11) (Bairoch 2004) using BLASTP. We used InterPro databases v32.0 (Zdobnov and Apweiler 2001; Quevillon *et al*. 2005) including Pfam, PRINTS, PROSITE, ProDom, and SMART to identify protein motifs and domains. Gene Ontology terms were obtained from the corresponding InterPro entries.

#### BUSCO assessment of annotation completeness

Annotated gene set completeness in terms of expected gene content was quantified using the BUSCO assessment tool (Waterhouse *et al*. 2018) for *N. melanderi* and seven other Apoidea species. Gene sets were first filtered to select the single longest protein sequence for any genes with annotated alternative transcripts. Gene set completeness assessments employed BUSCOv3.0.3 with HMMER 3.1b2 (Finn *et al*. 2011), and BLAST+ 2.7.1 (Camacho *et al*. 2009), using both the hymenoptera_odb9 and the insecta_odb9 BUSCO lineage datasets.

#### Transcription factor motif scans

We generated binding scores for 223 representative transcription factor (TF) binding motifs in the *N. melanderi* genome. Motifs representative of TF clusters with at least one ortholog in bees (Kapheim *et al*. 2015) were selected from FlyFactorSurvey (Zhu *et al*. 2011). After masking tandem repeats with Tandem Repeat Finder, we produced normalized genome-wide scoring profiles for each selected TF motif in the genome based on sliding windows of 500 bp with 250 bp overlap. We used the HMM-based motif scoring program Stubb (Sinha *et al*. 2006) with a fixed transition probability of 0.0025 and a background state nucleotide distribution learned from 5 kb regions without coding features of length > 22 kb. We then normalized these motif scores using two different methods. First, we created a “Rank Normalized” matrix, to normalize the window scores across each motif on a scale of 0 (best) to 1 (worst). Second, we created a “G/C Normalized” matrix, by considering each window’s GC content. Motifs with high GC content are likely to produce a high Stubb score in a GC rich window. We thus separated genomic windows into 20 bins of equal size based on GC content, and performed rank-normalization separately within each bin. We next summarized motif scores at the gene level. For each gene, we calculated a score for each motif as *Pgm* = *1*-*(1*-*Ngm)*^*Wg*, where *Ngm* is the best normalized score for motif *m* among the *Wg* windows that fall within the regulatory region of the gene *g*. We defined the regulatory region of the gene in five different ways: *5Kup2Kdown* − 5000 bp upstream to 2000 bp downstream of a gene’s transcription start site (TSS), *5Kup* − 5000 bp upstream of a gene’s TSS, *1Kup* − 1000 bp upstream of a gene’s TSS, *NearStartSite* − all genomic windows that are closer to the gene’s TSS than any other gene TSS, *GeneTerr* − all genomic windows between the boundary positions of the nearest non-overlapping gene neighbors within at least 5000 bp upstream of the TSS.

We used the results of these target motif scans to check for transcription factor motif enrichment among gene sets of interest (i.e., genes under selection). For each normalization method and regulatory region, we created two motif target gene sets: a “conservative” set that contains only the top 100 genes by normalized score and a more “liberal” set that contains the 800 top genes. Enrichment tests for genes of interest were performed using the one-sided Fisher exact test for each of 1784 motif target sets defined using the two thresholds, both “G/C” and “Rank” normalization procedures, the *1Kup* (likely the core promoter) and *GeneTerr* (likely containing distal enhancers) regulatory region definitions, and each of the representative 223 motifs. Multiple hypothesis test corrections were performed using the Benjamini-Hochberg procedure (Benjamini and Hochberg 1995). For significantly enriched motifs (adjusted-p < 6E-04), we determined if an ortholog of the fly transcription factor protein was present in the *N. melanderi* genome using blastp with e-value < 10e-3 and *%* identity ≥ 50.

#### Transposable element identification

We performed a more detailed *de novo* investigation of transposable elements in the *N. melanderi* genome using raw sequencing reads in a genome assembly-independent approach. First, we filtered a subset of five million raw reads for mitochondrial contamination to avoid biasing the detection of highly repetitive sequences. This involved aligning reads to the genome assembly with bwa-mem (Li 2013), and evaluating read depth with bedtools (Quinlan and Hall 2010). We identified contigs and scaffolds with high coverage (≥ 500x) as potential mitochondrial sequences, based on the assumption that the number of sequenced mitochondrial copies is much higher than that of the nuclear genome.

These contigs and scaffolds were further analyzed for sequence similarity (blastn v. 2.2.28+) to the mitogenome of the closest available bee species, *Halictus rubicundus* (KT164656.1). We identified five scaffolds as putatively mitochondrial (scaffold235256, scaffold241193, scaffold252191, scaffold252994, scaffold257806). Reads aligning to these scaffolds were filtered from the analysis.

The remaining reads were used for repeat analysis in five iterations of the transposable element discovery program DnaPipeTE v1.1 (Goubert *et al*. 2015), following Stolle et al. (2018). Each iteration used a new set of the same number of reads randomly sampled from the filtered reads. The analysis was repeated for different number of reads to represent a genome sequence assembly length coverage of 0.20x-0.40x in steps of 0.05x. This series of repeat content estimates determines the amount of input data that provides a stable estimate of genomic repeat content, and thus ensures that adequate coverage has been obtained for accurate estimates. The final set of repetitive elements was generated based on 0.30x coverage, using RepeatMasker v4.0.7 and a 10% sequence divergence cut-off. Overlap between repetitive element annotations and genes was detected with bedtools.

### Orthology delineation

Orthologous groups (OGs) delineated across 116 insect species were retrieved from OrthoDB v9.1 (Zdobnov *et al*. 2017) to identify orthologs. The OrthoDB orthology delineation procedure employs all-against-all protein sequence alignments to identify all best reciprocal hits (BRHs) between genes from each pair of species. It then uses a graph-based approach that starts with BRH triangulation to build OGs containing all genes descended from a single gene in the last common ancestor of the considered species. The annotated proteins from the genomes of *N. melanderi* were first filtered to select one protein-coding transcript per gene and then mapped to OrthoDB v9.1 at the Insecta level, using all 116 species and an unpublished halictid bee genome (*Megalopta genalis*; Kapheim et al. unpublished) for orthology mapping. The OrthoDB orthology mapping approach uses the same BRH-based procedure as for building OGs, but only allowing proteins from the mapped species to join existing OGs.

### Phylogenomic analysis

We reconstructed a molecular species phylogeny from 2,025 universal single-copy orthologs among the protein sequences of 15 insects including *N. melanderi* (Table S1-S2). The protein sequences from each orthogroup were first aligned with Muscle 3.8.31 (Edgar 2004), then trimmed to retain only confidently aligned regions with TrimAl v1.3 (Capella-Gutierrez *et al*. 2009), and then concatenated to form the 15 species superalignment of 688,354 columns. The maximum likelihood phylogeny was then estimated using RAxML 8.0.0 (Stamatakis 2014), with the PROTGAMMAJTT substitution model, setting the body louse (*Pediculus humanus*) as the outgroup species, and performing 100 bootstrap samples to obtain support values.

With these data, we performed a comparative orthology analysis to identify genes with universal, widely shared, or lineage-specific/restricted distributions across the selected species, or with identifiable orthologs from other insect species from OrthoDB v9.1. Ortholog presence, absence, and copy numbers were assessed for all OGs across the 15 species to classify genes according to their orthology profiles. The categories (each mutually exclusive) included: 1) Single-copy in all 15 insect species; 2) Present in all 15 insect species; 3) Halictidae: Present in >=2 Halictidae but none of the other 11 species; 4) Apidae + Mrot: Present in >= 2 Apidae and *Megachile rotundata* but none of the other 11 species; 5) Apoidea: Present in >= 1 Halictidae, present in >= 1 Apidae and *Megachile rotundata* but none of the other 7 species; 6) Formicoidea: Present in >= 2 Formicoidea but none of the other 11 species; 7) Apoidea + Formicoidea: Present in >=2 Apoidea, present in >=1 Formicoidea but not in *Polistes dominula* or *Cephus cinctus* or *P. humanus*; 8) Pdom + Ccin: Present in *P. dominula* and *C. cinctus* but none of the other 13 species; 9) Pdom or Ccin or Phum: Present in >=2 of *P. dominula* or *C. cinctus* or *P. humanus* and none of the other 12 species; 10) Hymenoptera: Present in >=2 Hymenoptera and absent from *P. humanus*; 11) Present < 13: Present in <13 of the 15 species, i.e., a patchy distribution not represented by any other category; 12) Other orthology: Present in any other insect from OrthoDB v9.1; 13) No orthology: No identifiable orthology at the OrthoDB v9.1 Insecta level.

### Population genetic analysis

#### SNP discovery and filtering

We used sequences generated from the 18 females and one male to characterize genetic variants following GATK best practices (https://software.broadinstitute.org/gatk/best-practices/). Reads were pre-processed by quality trimming using sickle with default parameters (Joshi and Fass 2011). We then converted paired reads to BAM format and marked adapters with Picard tools (“Picard.

http://picard.sourceforge.net/. Accessed 12 January 2016”). Reads were aligned to the genome with bwa-mem wrapped through Picard tools (CLIPPING_ATTRIBUTE=XT, CLIPPING_ACTION=2, INTERLEAVE=true, NON_PF=true). Alignments were then merged with MergeBamAlignment (CLIP_ADAPTERS=false, CLIP_OVERLAPPING_READS=true, INCLUDE_SECONDARY_ALIGNMENTS=true, MAX_INSERTIONS_OR_DELETIONS=-1, PRIMARY_ALIGNMENT_STRATEGY=MostDistant, ATTRIBUTES_TO_RETAIN=XS). PCR duplicates were marked with the function MarkDuplicatesWithMateCigar (OPTICAL_DUPLICATE_PIXEL_DISTANCE=2500, MINIMUM_DISTANCE=300). We next identified and realigned around indels using the Picard tools functions RealignerTargetCreator and IndelRealigner.

We performed variant calling in two rounds. The first pass was to generate a high quality SNP set that could be used for base quality recalibration, followed by a second pass of variant calling. For both rounds, we used the HaplotypeCaller function in Picard tools (-- variant_index_type LINEAR, --variant_index_parameter 128000, -ERC GVCF), followed by joint genotyping for the 18 females and individual genotyping for the male sample (GenotypeGVCFs). Haplotype caller was run set with μloidy level = 2n for all samples, including the haploid male. The latter was used to identify low-confidence or spurious SNPs that could be filtered from the female calls.

Variant filtering followed the GATK generic recommendations (--filterExpression “QD < 2.0, FS > 60.0, MQ < 40.0, ReadPosRankSum < -8.0, --restrictAllelesTo BIALLELIC). These were further filtered for SNPs identified as heterozygous in the male sample and for which genotypes were missing in any sample (--max-missing-count 0).

This set of high-confidence SNPs was used as input for base quality score recalibration for the 18 females. The second round of variant calling and filtering for these samples followed that of the first round, with the exception that we allowed missing genotypes in up to 8 samples. We then applied a final, more stringent set of filters using vcftools (Danecek *et al*. 2011) (--min-meanDP 5, --max-missing-count 4, --maf 0.05, --minGQ 9, --minDP 3). This yielded a final set of 412,800 high confidence SNPs used in the downstream analyses (File S1).

#### Structure analysis

We evaluated the potential for population structure by estimating heterozygosity, relatedness, and Hardy-Weinberg disequilibrium within our samples using vcftools. We also used ADMIXTURE v.1.3 (Alexander *et al*. 2009) to look for evidence of population structure (N=18 diploids). We randomly extracted SNPS that were at least 1000bp apart across the genome and ran K = 1-4 for three independent datasets.

#### SNP function

We identified the functional role (e.g., upstream, synonymous, non-synonymous, etc.) of SNPs using SNPEFF (Cingolani *et al*. 2012) for all SNPs within our data set (N = 412,800).

#### Genetic diversity

We characterized genetic diversity by evaluating pi and Tajima’s D in 10Kb and 1Kb windows with vcftools (--window-pi, --TajimaD, --site-pi). We mapped gene models to these windows with bedtools intersect, and Tajima’s D and pi values were averaged over each gene model using the aggregate function in R (Team 2016). We then calculated the cumulative percentile for pi and Tajima’s D for each gene using the ecdf function in R. These percentiles were then multiplied and recalculated. Genes for the joint percentile of pi and Tajima’s D that fell in the lowest 5% were considered to be under ongoing positive selection. To estimate genetic diversity across the genome in windows, we first calculated coverage at each site within 1Kb windows across the genome using bedtools coverage. Within each window, we estimated the proportion of sites with at least 5 reads of coverage. We used this value as the denominator to calculate pi within 1Kb windows.

#### Effective population size and demography

We estimated Ne using SMC++ (Terhorst *et al*. 2017). We randomly selected 4 large scaffolds (> 1 Kb) and estimated effective population size of our single *Nomia melanderi* population from 1000 to 100000 years before present. We assumed a single generation per year and a mutation rate of 6.8x10^−9^ (Liu *et al*. 2017). For each scaffold, we created 6 datasets by randomly selecting between 5 and 8 individuals without replacement. We used these files to estimate Ne using the cross-validation for each scaffold.

We evaluated the possibility of recent demographic changes by estimating Tajima’s D in 1000bp windows across the genome for all samples (Tajima 1989).

### Evolutionary rate analysis

Single copy orthologs were extracted from OGs identified above for *Lasioglossum albipes, Dufourea novaengliae, M. genalis,* and *N. melanderi*. Peptide alignments were obtained by running GUIDANCE2 (Penn *et al*. 2010) with the PRANK aligner (Löytynoja 2014) and species tree ((Dnov:67.51,(Nmel:58.18,(Mgen:47.03,Lalb:47.03):11.15):9.33); (Branstetter *et al*. 2017)) on each orthogroup. Low scoring residues (scores < 0.5) were masked to N using GUIDANCE2 to mask poor quality regions of each alignment. PAL2NAL (Suyama *et al*. 2006) was used to back-translate aligned peptide sequences to CDS and format alignments for PAML. PAML (Yang 2007) was run to evaluate the likelihood of multiple hypothesized branch models of dN/dS relative to two null models with trees and parameters as follows:

M0: (Dnov:67.51,(Nmel:58.18,(Mgen:47.03,Lalb:47.03):11.15):9.33); (model = 0, fix_omega = 0, omega = 0.2; all branches same omega)

M1a: (Dnov:67.51,(Nmel:58.18 #1, (Mgen:47.03,Lalb:47.03):11.15):9.33);

(model = 2, fix_omega = 1, omega = 1; neutral evolution for Nmel branch)

M2a: (Dnov:67.51,(Nmel:58.18 #1,(Mgen:47.03,Lalb:47.03):11.15):9.33); (model = 2, fix_omega = 0, omega = 0.2; Nmel branch different omega)

M1b: (Dnov:67.51,(Nmel:58.18,(Mgen:47.03 #1,Lalb:47.03):11.15):9.33); (model=2, fix_omega=1, omega = 1; neutral evolution for Mgen branch)

M2b: (Dnov:67.51,(Nmel:58.18,(Mgen:47.03 #1,Lalb:47.03):11.15):9.33); (model=2, fix_omega=0, omega=0.2; Mgen branch different omega)

M1c: (Dnov:67.51,(Nmel:58.18,(Mgen:47.03,Lalb:47.03 #1):11.15):9.33); (model=2, fix_omega=1, omega=1; neutral evolution for Lalb branch)

M2c: (Dnov:67.51,(Nmel:58.18,(Mgen:47.03,Lalb:47.03 #1):11.15):9.33); (model=2, fix_omega=0, omega=0.2; Lalb branch different omega)

Orthogroups with dS>2 were removed, and likelihood ratio tests were performed to determine the most likely value of omega for each branch.

### Functional Enrichment Tests

We performed all tests of functional enrichment using the GOstats package (Gentleman and Falcon 2013) in R version 3.4.4. We used terms that were significantly enriched (p < 0.05) to build word clouds with the R packages tm (Feinerer *et al*. 2008), SnowballC (Bouchet-Valat 2014), and wordcloud (Fellows 2018).

### Data Availability

Sequence data are available at NCBI (BioProject PRJNA495036). The genome assembly is available at NCBI (BioProject PRJNA494873). Genetic variants and genotypes are available in VCF format in File S1. TF binding motif scores are in File S2. Repetitive DNA content is in File S3. SNP effects are in File S4. The genome annotation (GFF format) is in File S5. All supplementary tables (Table S1-S8) and files (Files S1-S5) have been deposited at FigShare.

## Results and Discussion

The *N. melanderi* genome assembly resulted in 268,376 scaffolds (3,194 > 1 kb) with an N50 scaffold length of 2.05 Mb (Table 1). Total size is estimated to be 299.6 Mb, based on a k-mer analysis with k = 17 and a peak depth of 70. CEGMA analysis indicated 244 of 248 (98.39%) core eukaryotic genes were completely assembled, and 10.25% of the detected CEGMAs had more than one ortholog. BUSCO analyses indicated 98.8% of Insecta BUSCOs were complete in the assembly (Table S3).

**Table 1.** Comparison of genome assemblies among bees, including *Nomia melanderi*.

| Species | Genome size (Mb) | Number scaffolds | N50 Scaffold length | Predicted Genes | Coverage (X) | Reference |
| --- | --- | --- | --- | --- | --- | --- |
| *Nomia melanderi* | 299.6 | 268,376<br>(3,194 > 1kb) | 2,054,768 | 10,847 | 75 | --- |
| *Lasioglossum albipes* | 416 | 41,377 | 616,426 | 13,448 | 96 | (Kocher *et al.* 2013) |
| *Dufourea novaeangliae* | 291 | 84,187 | 2,397,596 | 12,453 | 133 | (Kapheim *et al.* 2015) |
| *Megachile rotundata* | 273 | 6,266 | 1,699,680 | 12,770 | 272 | (Kapheim *et al.* 2015) |
| *Bombus impatiens* | 248 | 5,559 | 1,399,493 | 15,896 | 108 | (Sadd *et al.* 2015) |

Our official gene set includes 10,847 predicted protein-coding gene models. This is likely to be a relatively complete gene set, as 96.0% of Insecta BUSCOs were identified as complete, which is comparable to other bee genomes (Table S3). Most (8,075) of the predicted genes belong to ancient OGs that include orthologs in vertebrate species. However, there were 819 genes without any known orthologs (Fig. 1B). Our comparative analysis with representative Hymenoptera species and the outgroup, *P. humanus*, identified 2,025 single-copy orthologs from which we constructed the molecular species phylogeny that confidently μlaces Halictidae as a sister group to the combined Apidae and Megachilidae groups within Apoidea (Fig. 1C). Orthology delineation showed that 92.2% of *N. melanderi* predicted genes have orthologs in other insects and only 16 of them were unique to the family Halictidae (Fig. 1C). Transcription factor motif binding scores for each gene are available in File S2.

In a genome-assembly independent approach using short reads and DnaPipeTE, we assembled 54,236 repetitive elements, suggesting that 37.5% of the *N. melanderi* genome is repetitive content (File S3; Fig. 1D). We identified transposable elements from all major groups (LTR, LINE, SINE, DNA, Helitron) and other elements with similarities to unclassified repeats (7,866 total annotated repeats), but unknown elements are the most abundant type of transposon (25.5%) (Fig. 1D), showing no similarities to known repetitive elements, conserved domains, or sequences in NCBI’s non-redundant nt database.

Of annotated transposable elements, LINE retrotransposons (most common: I and Jockey) were the most abundant, followed by LTR retrotransposons (most common: Gypsy) and small amounts of DNA (mostly Tc1-Mariner, PiggyBac, hAT and Kolobok) or other transposons (File S3). Some annotations suggest the presence of Crypton, Helitron and Maverick elements as well as 5S/tRNA SINE (File S3). A majority of the detected retroelements show little sequence divergence, indicating recent activity, particularly Gypsy (LTR), Copia (LTR), I (LINE) and R2 (LINE).

Annotation of the genome assembly yielded 25.93 Mbp of masked sequences (8.59% at 10% sequence divergence), which is less than the repetitive fraction of >37% inferred by DnaPipeTE. Even at a 20% sequence divergence threshold, only 43.36 Mbp (14.37%) were masked, suggesting that a substantial fraction of the repetitive part of the genome is not part of the genome assembly, likely due to the technical limitations in assembling repetitive elements from short reads.

Our population genetic analysis indicated our population is panmictic. We did not find any evidence of population structure among our samples. Across all three datasets run through STRUCTURE, the lowest CV error was found for K = 1 (CV = 0.68) (Fig. 2A). Likewise, pairwise relatedness estimates based on the unadjusted λ_*i*_k statistic were close to 0 (-0.084 - - 0.047) for all females in our population (Yang *et al*. 2010).

**Figure 2.**
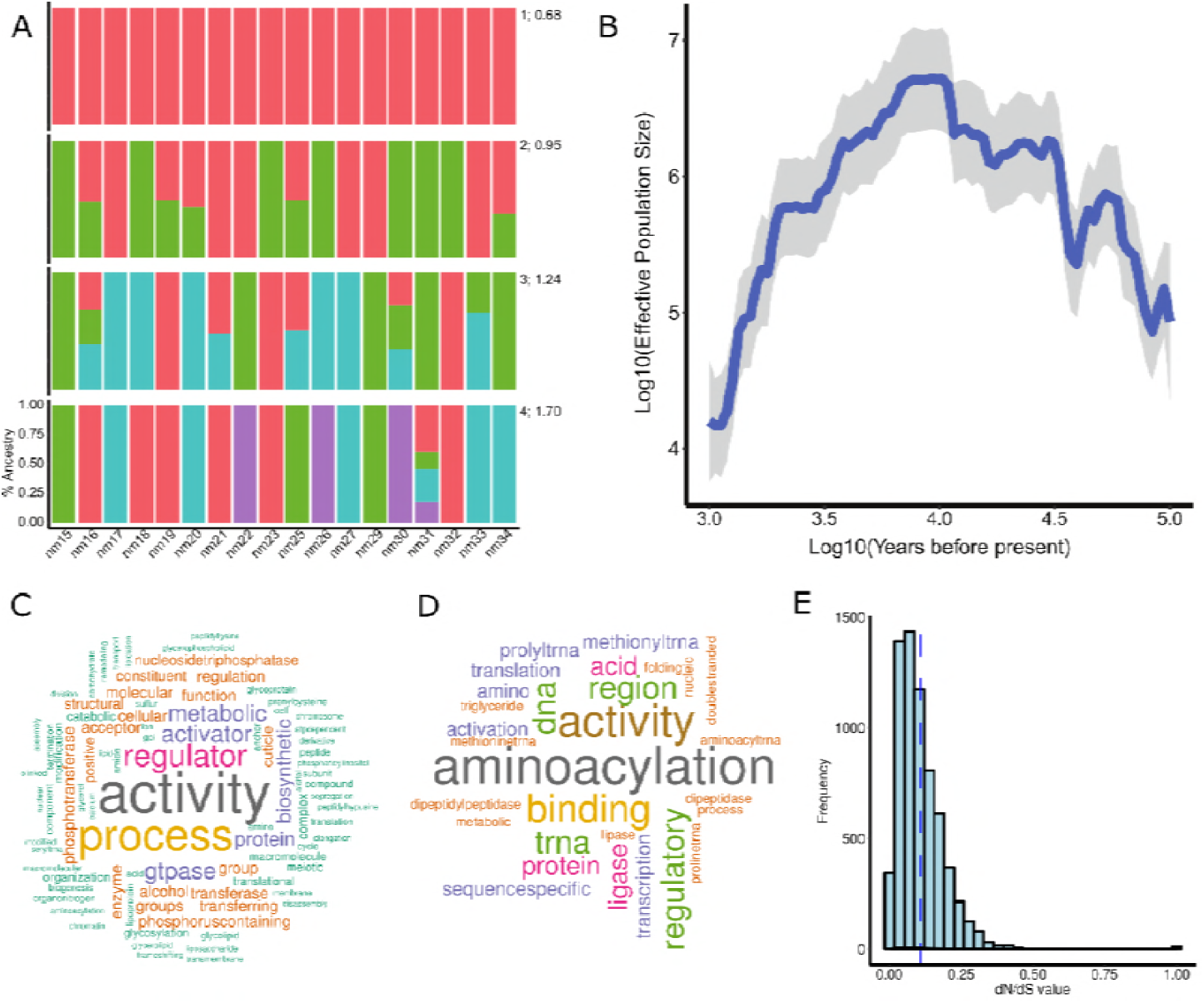
*N. melanderi* population genetics. (A) Samples most likely originate from a single source population. We tested for population structure for K=1-4 (right numbers) and found that the most likely K = 1 (average CV error = 0.68 across three independent runs). K:CV is given to the right of each row. (B) Estimates of N_e_ show evidence for a decline in effective population size in our alkali bee population, beginning about 10,000 years before present. Blue line, median estimated N_e_; shaded gray area, 95% confidence intervals. (C) Genes under positive selection are significantly enriched for molecular functions and biological processes related to tRNA transfer and binding. (D) Genes with a slower evolutionary rate (dN/dS) in *N. melanderi* than in other halictid bees are significantly enriched for processes and functions related to transcription and translation. In B and C, the size of the word corresponds to the frequency to which that term appears on a list of significantly enriched GO terms. (E) The distribution of dN/dS values for *N. melanderi* genes are skewed toward zero, and none are greater than 1. Blue dashed line, mean dN/dS.

Solitary bees are expected to have high genetic diversity and large effective population sizes (Romiguier *et al*. 2014), and recent census data suggests there are 17 million females nesting in our study population in the Touchet Valley (Washington, USA) (Cane 2008). However, we find several lines of evidence to suggest that effective population size of our *N. melanderi* population has declined in the recent past. First, our estimates of genetic diversity were surprisingly low. Three of the 18 females in our dataset had significantly higher homozygosity than expected (p < 0.05). Genetic diversity (pi) across the genome in 1Kb windows (corrected for coverage, see Methods) was estimated to be 0.00153. This is intermediate to diversity previously estimated for *Apis mellifera* (0.0131, (Harpur *et al*. 2014)) and *Bombus impatiens* (0.002, (Harpur *et al*. 2017)).

Second, the genome-wide average Tajima’s D was significantly greater than 0 (one-way T-test; mean = 0.77 +/- 0.002 SE; p < 0.00001) indicating a recent population decline.

Third, N_e_ is predicted to have declined within the last 10,000 years (Fig. 2B). In the last 2,000 years, N_e_ has had a median of 12,554 individuals (range: 3,119-3,978,942). The long, slow population decline reflected in our samples corresponds to a period during which much of Washington state was underwater due to glacial flooding, known as the Missoula Floods. Our study area, Touchet Valley, was under Lake Lewis during this time, and was thus uninhabitable for ground-nesting bees.

More recent fluctuations in N_e_ may reflect less catastrophic events. Seed growers have maintained large nesting areas (“bee beds”) for alkali bees within a 240 km^2^ watershed that encompasses our sampling area for several decades (Cane 2008). Some of these bee beds are among the largest nesting aggregations ever recorded, at up to 278 nests per m^2^ However, survey data suggests there are large fluctuations in population size, as the population increased 9-fold over an eight year period (1999-2006) (Cane 2008). Records from individual bee beds reflect these fluctuations. For example, a bed that was started in 1973 grew from 550 nesting females to 5.3 million nesting females in 33 years (Johansen *et al*. 1978; Cane 2008). However, other beds were destroyed or abandoned for decades at a time, only to be recolonized later. A large population crash occurred in the 1990s, likely due to use of a new pesticide (Cane 2008), and flooding events have caused massive valley-wide reproductive failures (Stephen 2003). Our wide range of N_e_ estimates and signatures of genetic bottlenecks likely reflect these population fluctuations.

Our selection scan revealed 479 *N. melanderi* genes under positive directional selection. Genes under selection were highly conserved, and the age distribution was similar to the distribution across all predicted genes (χ^2^ = 54, *d*.f. = 48, p = 0.26; Table S4). Genes showing signatures of ongoing positive selection were enriched for functions related to tRNA transfer and DNA/nucleosome binding (Fig. 2C, Table S5). Because DNA binding is typically an indicator of transcription factor activity, we performed enrichment analysis of genes under selection with our previously defined transcription factor motif target sets (File S2). The most enriched motif target sets (adjusted-p < 6E-04) included transcription factors involved in neural differentiation (*brick*-*a*-*brack 1*, *prospero*, *nubbin*, *zelda*, *twin*-*of*-*eyeless*, *pox meso*, *worniu*) and neural secretory functions (*dimmed*) (Table S6). We identified 505,203 functional predictions for 412,800 variable sites (SNPs) within 9,692 genes, most of which are intergenic (File S4).

Our analysis of evolutionary rates included 6,644 single-copy orthologs, most of which (95%) were evolving at similar rates across all four halictid bee lineages. We identified 61 *N. melanderi* genes that are evolving at a significantly different rate from other halictid bees (Table S7). Of these, the majority (74%) are evolving slower than in other lineages. These genes are significantly enriched for functions related to transcription and translation (Fig. 2D, Table S8). The distribution of estimated dN/dS values for *N. melanderi* genes was skewed toward zero, with a notable absence of values greater than one (Fig. 2E). This suggests that most genes in our analysis show evidence of neutral or purifying selection. This result is likely influenced by the vast evolutionary distance separating the four halictid lineages, which shared a common ancestor > 150 million years ago (Branstetter *et al*. 2017). Our set of single-copy orthologs was thus limited to highly conserved genes.

In conclusion, we present a high quality draft genome assembly of the solitary alkali bee, *N. melanderi*, that will be a valuable resource for both basic and applied research communities.

## Acknowledgements

We are grateful to M. Ingham, M. Buckley, and M. Wagoner for allowing us to collect from their bee beds. J. Dodd and Forage Genetics International provided lab space while in the field. Sequencing was performed by the Roy J. Carver Biotechnology Center at University of Illinois at Urbana-Champaign (UIUC). Computational support was provided by University of Utah Center for High Performance Computing and UIUC CNRG/Biocluster. J. Johnson (Life Sciences Studios) created the illustration in Fig. 1. Funding was provided by Utah Agricultural Experiment Station project 1297 (KMK) and grants from the USDA-ARS Alfalfa Pollinator Research Initiative (KMK) and USDA-NIFA grant # 2018-67014-27542 (KMK). Additional funding was provided by a Smithsonian Institution Competitive Grants Program for Biogenomics (WTW, KMK, BMJ), Swiss National Science Foundation grant PP00P3_170664 (RMW), and general research funds from the Smithsonian Tropical Research Institute (WTW).

## Supplementary Files

**Table S1. Selected species for orthology analysis.** All protein sets were collated from OrthoDB v9.1, except the new genome assembly presented here (*Nomia melanderi*) and the unpublished *Megalopta genalis* genome assembly (Kapheim et al., unpublished; BioProject PRJNA494872).

**Table S2. Comparison of the eight selected Apoidea genomes used for orthology and phylogenomics analyses, as well as BUSCO assessments.**

**Table S3. BUSCO genome and gene set assessments of selected Hymenoptera.** Complete BUSCOs (C); Complete and single-copy BUSCOs (S); Complete and duplicated BUSCOs (D); Fragmented BUSCOs (F); Missing BUSCOs (M); Total BUSCO groups searched (*n*) for Insecta (Ins.) and Hymenoptera (Hym.) assessment sets.

**Table S4. Genes under selection.** This is a list of *N. melanderi* genes found to undergoing positive selection via population genetic analyses. Orthogroup IDs and orthogroup ages are also included.

**Table S5. GO enrichment for genes under selection.** This is the GOstats output for enrichment tests of the set of genes under positive selection in *N. melanderi*.

**Table S6. Transcription factor motif enrichments among genes under positive selection in *N. melanderi*.**

**Table S7. Model likelihood scores and predicted omega values from PAML.** Omega values and model likelihood results from PAML analysis of 6676 orthogroups. Omega values are given for *N. melanderi* for best-fitting model(s), as well as additional foreground and background branches. Raw and FDR corrected p-values for each model likelihood test are also provided. Orthogroups with dS>2 in at least one branch in the best-fitting model were filtered from the results presented in the main text.

**Table S8. GO enrichment for PAML results.** This is the GOstats output for functional enrichment tests on the set of genes identified as evolving faster or slower in *N. melanderi*, as compared to other species in Halictidae.

**File S1. VCF file.** File containing genetic variant information for the 18 females included in the population genetic analysis.

**File S2. Collection of normalized TF motif binding score matrices.** The ten files in this archive are matrices of gene-level motif binding scores for *N. melanderi* for each combination of the two different normalization procedures (“rank” and “gc”) and the five different regulatory region definitions. The rows are genes, the columns motifs, and the scores are the length-adjusted normalized scores which range from 0 (best) to 1 (worst). Scores of “2” signal a missing motif score, likely indicating that the regulatory region was small and masked by tandem repeats. More details and datasets for additional bee species are available at http://veda.cs.uiuc.edu/beeMotifScores/.

**File S3. Repetitive Elements in the genome of Nomia melanderi (Nme).** This contains supplementary information on the repetitive and transposable elements in Nme, including additional figures (S3.Nme.Repeats.report.pdf), tables (Nme.basepairs.by.type.txt: Nme basepairs per repeat type; Nme.TE.groups.counts.txt: Nme counts per TE group; Nme.TE.elements.counts.txt: Nme counts per TE elements, Nme.TE.RM.annotation.report.txt: Nme repeats annotated with RepeatMasker), and annotation files (Nme.TE.fa: Nme repetitive elements fasta sequences; Nme.TE.gff: Nme sequence assembly annotation of repetitive elements (gff)).

**File S4. Predicted SNP effects.** This file has two tabs describing the predicted effects of each filtered SNP. The ‘functionalPredictions’ tab indicates the number of SNPs found within each protein-coding gene, and its position/effect relative to the gene structure. The tab ‘predictedEffects’ describes the location (Scaffold, Position) of each filtered SNP associated with a protein coding gene. The reference and alternate alleles are provided, along with the gene name it is associated with.

**File S5. Gene annotation file (GFF) for *N. melanderi*.**

## References

Alexander, D. H., J. Novembre, and K. Lange, 2009 Fast model-based estimation of ancestry in unrelated individuals. Genome Res. 19: 1655–64.

Bairoch, A., 2004 Swiss-Prot: Juggling between evolution and stability. Brief. Bioinform. 5: 39–55.

Batra, S. W. T., 1970 Behavior of alkali bee, *Nomia-melanderi*, within nest (Hymenoptera-Halictidae). Ann. Entomol. Soc. Am. 63: 400–406.

Batra, S. W. T., 1976 Comparative efficiency of alfalfa pollination by *Nomia melanderi, Megachile rotundata, Anthidium florentinum* and *Pithitis smaragdula* (Hymenoptera: Apoidea). Source J. Kansas Entomol. Soc. 49: 18–22.

Batra, S. W. T., 1972 Some properties of the nest-building secretions of *Nomia, Anthophora, Hylaeus* and other bees. J. Kansas Entomol. Soc. 45: 208–218.

Batra, S. W., and G. E. Bohart, 1969 Alkali bees: response of adults to pathogenic fungi in brood cells. Science 165: 607.

Benjamini, Y., and Y. Hochberg, 1995 Controlling the false discovery rate: a practical and powerful approach to multiple testing. J R Stat Soc Ser B 57:.

Benson, G., 1999 Tandem repeats finder: a program to analyze DNA sequences. Nucleic Acids Res 27: 573–580.

Birney, E., M. Clamp, and R. Durbin, 2004 GeneWise and GenomeWise. Genome Res 14: 988–995.

Bouchet-Valat, M., 2014 Snowball stemmers based on the C libstemmer UTF-8 library [R package SnowballC version 0.5.1].

Brady, S. G., S. Sipes, A. Pearson, and B. N. Danforth, 2006 Recent and simultaneous origins of eusociality in halictid bees. Proc. R. Soc. B Biol. Sci. 273: 1643–1649.

Branstetter, M. G., A. K. Childers, D. Cox-Foster, K. R. Hopper, K. M. Kapheim et al., 2018 Genomes of the Hymenoptera. Curr. Opin. Insect Sci. 25: 65–75.

Branstetter, M. G., B. N. Danforth, J. P. Pitts, B. C. Faircloth, P. S. Ward et al., 2017 Phylogenomic insights into the evolution of stinging wasps and the origins of ants and bees. Curr. Biol. 27: 1019–1025.

Camacho, C., G. Coulouris, V. Avagyan, N. Ma, J. Papadopoulos et al., 2009 BLAST+: architecture and applications. BMC Bioinformatics 10: 421.

Cane, J. H., 2008 A native ground-nesting bee (*Nomia melanderi*) sustainably managed to pollinate alfalfa across an intensively agricultural landscape. Apidologie 39: 315–323.

Cane, J. H., 2002 Pollinating bees (Hymenoptera: Apiformes) of U.S. alfalfa compared for rates of pod and seed set. J. Econ. Entomol. 95: 22–27.

Capella-Gutierrez, S., J. M. Silla-Martinez, and T. Gabaldon, 2009 trimAl: a tool for automated alignment trimming in large-scale phylogenetic analyses. Bioinformatics 25: 1972–1973.

Cingolani, P., A. Platts, L. L. Wang, M. Coon, T. Nguyen et al., 2012 A program for annotating and predicting the effects of single nucleotide polymorphisms, SnpEff: SNPs in the genome of *Drosophila melanogaster* strain w1118; iso-2; iso-3. Fly (Austin). 6: 80–92.

Danecek, P., A. Auton, G. Abecasis, C. A. Albers, E. Banks et al., 2011 The variant call format and VCFtools. Bioinformatics 27: 2156–2158.

Danforth, B. N., S. Cardinal, C. Praz, E. A. B. Almeida, and D. Michez, 2013 The impact of molecular data on our understanding of bee phylogeny and evolution. Annu. Rev. Entomol. 58: 57–78.

Danforth, B. N., C. Eardley, L. Packer, K. Walker, A. Pauly et al., 2008 Phylogeny of Halictidae with an emphasis on endemic African Halictinae. Apidologie 39: 86–101.

Van Deynze, A. E., S. Fitzpatrick, B. Hammon, M. H. McCaslin, D. H. Putnam et al., 2008 Gene flow in alfalfa: biology, mitigation, and potential impact on production.:

Edgar, R. C., 2004 MUSCLE: multiple sequence alignment with high accuracy and high throughput. Nucleic Acids Res 32: 1792–1797.

Edgar, R. C., and E. W. Myers, 2005 PILER: identification and classification of genomic repeats. Bioinformatics 21: i152–i158.

Elsik, C. G., K. C. Worley, A. K. Bennett, M. Beye, F. Camara et al., 2014 Finding the missing honey bee genes: Lessons learned from a genome upgrade. BMC Genomics 15: 1–29.

Feinerer, I., K. Hornik, and D. Meyer, 2008 Text mining infrastructure in R. J. Stat. Softw. 25: 1–54.

Fellows, I., 2018 wordcloud: Word Clouds. R package version 2.6.

Finn, R. D., J. Clements, and S. R. Eddy, 2011 HMMER web server: interactive sequence similarity searching. Nucleic Acids Res 39: W29–37.

Gentleman, R., and S. Falcon, 2013 Package “GOstats.”

Gibbs, J., S. G. Brady, K. Kanda, and B. N. Danforth, 2012 Phylogeny of halictine bees supports a shared origin of eusociality for *Halictus* and *Lasioglossum* (Apoidea: Anthophila: Halictidae). Mol. Phylogenet. Evol. 65: 926–939.

Goubert, C., L. Modolo, C. Vieira, C. ValienteMoro, P. Mavingui et al., 2015 De novo assembly and annotation of the Asian tiger mosquito (*Aedes albopictus*) repeatome with dnaPipeTE from raw genomic reads and comparative analysis with the yellow fever mosquito (*Aedes aegypti)*. Genome Biol. Evol. 7: 1192–1205.

Harpur, B. A., A. Dey, J. R. Albert, S. Patel, H. M. Hines et al., 2017 Queens and workers contribute differently to adaptive evolution in bumble bees and honey bees. Genome Biol. Evol. 9: 2395–2402.

Harpur, B. A., C. F. Kent, D. Molodtsova, J. M. Lebon, A. S. Alqarni et al., 2014 Population genomics of the honey bee reveals strong signatures of positive selection on worker traits. Proc Natl Acad Sci U S A 111: 2614–2619.

James, R. R., 2011 Bee importation, bee price data, and chalk-brood, in Western Alfalfa Seed Grower Association Winter Seed Conference, Las Vegas, NV.

Johansen, C. A., D. F. Mayer, and J. D. Eves, 1978 *Biology and management of the alkali bee*, Nomia melanderi *Cockrell (Hymenoptera: Halictidae)*. Washington State Entomology.

Johnson, A. D., R. E. Handsaker, S. L. Pulit, M. M. Nizzari, C. J. O’Donnell et al., 2008 SNAP: A web-based tool for identification and annotation of proxy SNPs using HapMap. Bioinformatics 24: 2938–2939.

Joshi, N., and J. Fass, 2011 Sickle: A sliding-window, adaptive, quality-based trimming tools for FastQ files.

Kapheim, K. M., 2017 Nutritional, endocrine, and social influences on reproductive physiology at the origins of social behavior. Curr. Opin. Insect Sci. 22: 62–70.

Kapheim, K. M., and M. M. Johnson, 2017a Juvenile hormone, but not nutrition or social cues, affects reproductive maturation in solitary alkali bees (*Nomia melanderi*). J. Exp. Biol. jeb.162255.

Kapheim, K. M., and M. M. Johnson, 2017b Support for the reproductive ground plan hypothesis in a solitary bee: Links between sucrose response and reproductive status. Proc. R. Soc. B Biol. Sci. 284: 2016–2406.

Kapheim, K. M., H. Pan, C. Li, S. L. Salzberg, D. Puiu et al., 2015 Social evolution. Genomic signatures of evolutionary transitions from solitary to group living. Science 348: 1139–1143.

Kocher, S. D., C. Li, W. Yang, H. Tan, S. V Yi et al., 2013 The draft genome of a socially polymorphic halictid bee, *Lasioglossum albipes*. Genome Biol. 14: R142.

Li, H., 2013 Aligning sequence reads, clone sequences and assembly contigs with BWA-MEM. ARXIV 1303.3997:

Liu, H., Y. Jia, X. Sun, D. Tian, L. D. Hurst et al., 2017 Direct determination of the mutation rate in the bumblebee reveals evidence for weak recombination-associated mutation and an approximate rate constancy in insects. Mol. Biol. Evol. 34: 119–130.

Löytynoja, A., 2014 Phylogeny-aware alignment with PRANK, pp. 155–170 in Multiple Sequence Alignment Methods. Methods in Molecular Biology (Methods and Protocols), edited by D. Russell. Humana Press, Totowa, NJ.

Penn, O., E. Privman, H. Ashkenazy, G. Landan, D. Graur et al., 2010 GUIDANCE: a web server for assessing alignment confidence scores. Nucleic Acids Res. 38: W23–8.

Picard. http://picard.sourceforge.net/. Accessed 12 January 2016.

Price, A. L., N. C. Jones, and P. A. Pevzner, 2005 De novo identification of repeat families in large genomes. Bioinformatics 21: i351–i358.

Quevillon, E., V. Silventoinen, S. Pillai, N. Harte, N. Mulder et al., 2005 InterProScan: protein domains identifier. Nucleic Acids Res 33: W116–20.

Quinlan, A. R., and I. M. Hall, 2010 BEDTools: a flexible suite of utilities for comparing genomic features. Bioinformatics 26: 841–842.

Rehan, S. M., and A. L. Toth, 2015 Climbing the social ladder: The molecular evolution of sociality. Trends Ecol. Evol. 30: 426–433.

Robinson, G. E., C. M. Grozinger, and C. W. Whitfield, 2005 Sociogenomics: Social life in molecular terms. Nat. Rev. Genet. 6: 257–270.

Romiguier, J., J. Lourenco, P. Gayral, N. Faivre, L. A. Weinert et al., 2014 Population genomics of eusocial insects: the costs of a vertebrate-like effective population size. J. Evol. Biol. 27: 593–603.

Sadd, B. M., S. M. Barribeau, G. Bloch, D. C. De Graaf, P. Dearden et al., 2015 The genomes of two key bumblebee species with primitive eusocial organization. Genome Biol. 16: 76.

Sinha, S., Y. Liang, and E. Siggia, 2006 Stubb: a program for discovery and analysis of cis - regulatory modules. Nucleic Acids Res. 34: W555–W559.

Smit AFA, Hubley R, Green P: RepeatMasker. Available at: http://www.repeatmasker.org. [Accessed 9 April 2013].

Stamatakis, A., 2014 RAxML version 8: a tool for phylogenetic analysis and post-analysis of large phylogenies. Bioinformatics 30: 1312–3.

Stanke, M., O. Keller, I. Gunduz, A. Hayes, S. Waack et al., 2006 AUGUSTUS: A b initio prediction of alternative transcripts. Nucleic Acids Res. 34:.

Stephen, W. P., 2003 Solitary bees in North American agriculture: a perspective, pp. 41–66 in For Nonnative Crops, Whence Pollinators of the Future?, edited by K. Strickler and J. H. Cane. Entomol. Soc. Am., Lanham, MD.

Stolle, E., R. Pracana, P. Howard, C. I. Paris, S. J. Brown et al., 2018 Degenerative expansion of a young supergene. bioRxiv 326645.

Suyama, M., D. Torrents, and P. Bork, 2006 PAL2NAL: robust conversion of protein sequence alignments into the corresponding codon alignments. Nucleic Acids Res. 34: W609–12.

Tajima, F., 1989 The effect of change in population size on DNA polymorphism. Genetics 123:.

Team, R. C., 2016 R: A language and environment for statistical computing.

Terhorst, J., J. A. Kamm, and Y. S. Song, 2017 Robust and scalable inference of population history from hundreds of unphased whole genomes. Nat. Genet. 49: 303–309.

Trapnell, C., L. Pachter, and S. L. Salzberg, 2009 TopHat: Discovering splice junctions with RNA-Seq. Bioinformatics 25: 1105–1111.

Trapnell, C., A. Roberts, L. Goff, G. Pertea, D. Kim et al., 2012 Differential gene and transcript expression analysis of RNA-seq experiments with TopHat and Cufflinks. Nat. Protoc. 7: 562–578.

U.S. Department of Agriculture, N. A. S. S., 2014 Census of Agriculture Summary and State Data 2012: U.S. Government Printing Office.

Waterhouse, R. M., M. Seppey, F. A. Simão, M. Manni, P. Ioannidis et al., 2018 BUSCO Applications from quality assessments to gene prediction and phylogenomics. Mol. Biol. Evol. 35: 543–548.

Wcislo, W. T., and M. S. Engel, 1996 Social behavior and nest architecture of nomiine bees (Hymenoptera: Halictidae; Nomiinae). J. Kansas Entomol. Soc. 69: 158–167.

Wcislo, W. T., and J. H. Fewell, 2017 Sociality in bees, pp. 50–83 in Comparative Social Evolution, edited by D. R. Rubenstein and P. Abbot. Cambridge University Press, Cambridge.

Wilson, E. O., 1971 The Insect Societies. Harvard University Press, Cambridge, Massachusetts.

Xu, Z., and H. Wang, 2007 LTR_FINDER: an efficient tool for the prediction of full-length LTR retrotransposons. Nucleic Acids Res. 35: W265–W268.

Yang, Z., 2007 PAML 4: Phylogenetic analysis by maximum likelihood. Mol. Biol. Evol. 24: 1586–1591.

Yang, J., B. Benyamin, B. P. McEvoy, S. Gordon, A. K. Henders et al., 2010 Common SNPs explain a large proportion of heritability for human height. Nat. Genet. 42: 565–569.

Zdobnov, E. M., and R. Apweiler, 2001 InterProScan--an integration platform for the signature-recognition methods in InterPro. Bioinformatics 17: 847–848.

Zdobnov, E. M., F. Tegenfeldt, D. Kuznetsov, R. M. Waterhouse, F. A. Simão et al., 2017 OrthoDB v9.1: cataloging evolutionary and functional annotations for animal, fungal, plant, archaeal, bacterial and viral orthologs. Nucleic Acids Res. 45: D744–D749.

Zhu, L. J., R. G. Christensen, M. Kazemian, C. J. Hull, M. S. Enuameh et al., 2011 FlyFactorSurvey: A database of Drosophila transcription factor binding specificities determined using the bacterial one-hybrid system. Nucleic Acids Res. 39: 111–117.

